# Yet another de novo genome assembler

**DOI:** 10.1101/656306

**Authors:** Robert Vaser, Mile Šikić

## Abstract

Advances in sequencing technologies have pushed the limits of genome assemblies beyond imagination. The sheer amount of long read data that is being generated enables the assembly for even the largest and most complex organism for which efficient algorithms are needed. We present a new tool, called Ra, for de novo genome assembly of long uncorrected reads. It is a fast and memory friendly assembler based on sequence classification and assembly graphs, developed with large genomes in mind. It is freely available at https://github.com/lbcb-sci/ra.

This work has been supported in part by the Croatian Science Foundation under the project Single genome and metagenome assembly (IP-2018-01-5886), and in part by the European Regional Development Fund under the grant KK.01.1.1.01.0009 (DATACROSS). In addition, M.Š. is partly supported by funding from A*STAR, Singapore.

## I. INTRODUCTION

The pace of improvement in sequencing technologies is staggering. From modest fragments up to a thousand nucleotides obtained with the first two generations of sequencing, the read length increased manifold after just a few decades, but with a setback in accuracy. Nonetheless, the increase in maximal sequencing length facilitated significant advances in contiguity of genome assemblies. Both leaders of the third generation of sequencing, Oxford Nanopore Technologies (ONT) and Pacific Biosciences (PB), are continuously improving their methods and throughput. Their novel protocols enable the generation of ultra-long [1] or highly-accurate reads [2], which will mitigate the assembly problem even further. However, de novo assemblies for larger genomes are still fragmented and substantial amount of computational resources is needed to acquire them.

There is a vast amount of available genome assemblers today. Some are specialized for short reads, others for long, and the middle balances the advantages and disadvantages of both sides. Almost all employ graph based techniques to retrieve contiguous chains of sequenced reads, followed by a polishing step to get rid of the remaining sequencing errors. Short read assemblers are successors of the De Bruijn graph approach [3], while long read assemblers compute pairwise overlaps between all reads in order to build the string graph [4] or some of its variations, like the best overlap graph [5] and the assembly graph [6]. Portion of the research shifted to the generalization of de Bruijn graphs to make them more resilient to error-ridden third generation data. Presence or absence of read error-correction prior the assembly is another criterion by which the assemblers distinguish themselves amongst others. A list of available tools for long read assembly and their characteristics can be observed in Table 1.

**TABLE I.** TOOLS FOR DE NOVO GENOME ASSEMBLY OF LONG READS

| Tool | Approach | Reference |
| --- | --- | --- |
| Canu | Error correction → Bogart | [7] |
| Falcon | Error correction → String graph | [8] |
| Flye | ABruijn graph | [9] |
| Hinge | Repeat-aware BOG | [10] |
| Miniasm | Assembly graph | [6] |
| Wtdbg2 | Fuzzy Bruijn graph | [11] |

In this paper we present yet another approach to genome assembly of long reads, incorporated in a tool called Ra (short for <u>R</u>apid <u>A</u>ssembler), which is based on the Overlap-Layout-Consensus paradigm (OLC) and is a mixture of well established concepts developed in a memory friendly manner.

## II. METHODS

Ra uses pairwise overlaps generated by minimap2 [12] for a given set of raw sequences to build an assembly graph, a directed graph that is both Watson-Crick complete and containment free [6]. As a preprocessing step, it trims sequence adapters, purges chimeric sequences and removes false overlaps induced by repetitive genomic regions located at sequence ends. This is achieved by examining their pile-o-grams, which are produced directly from pairwise overlaps [10]. After graph construction, Ra follows the default graph simplification path, i.e. transitive reduction, tip removal and bubble popping. Leftover tangles are resolved by cutting short overlaps. Linear paths of the assembly graph are extracted and passed to the consensus module Racon [13] to iteratively increase the accuracy of the reconstructed genome. Further-more, if second generation sequencing data is available, it can be used to further increase the accuracy of the assembly. Pseudocode of the whole Ra pipeline can be seen in Algorithm 1.

**Algorithm 1.**
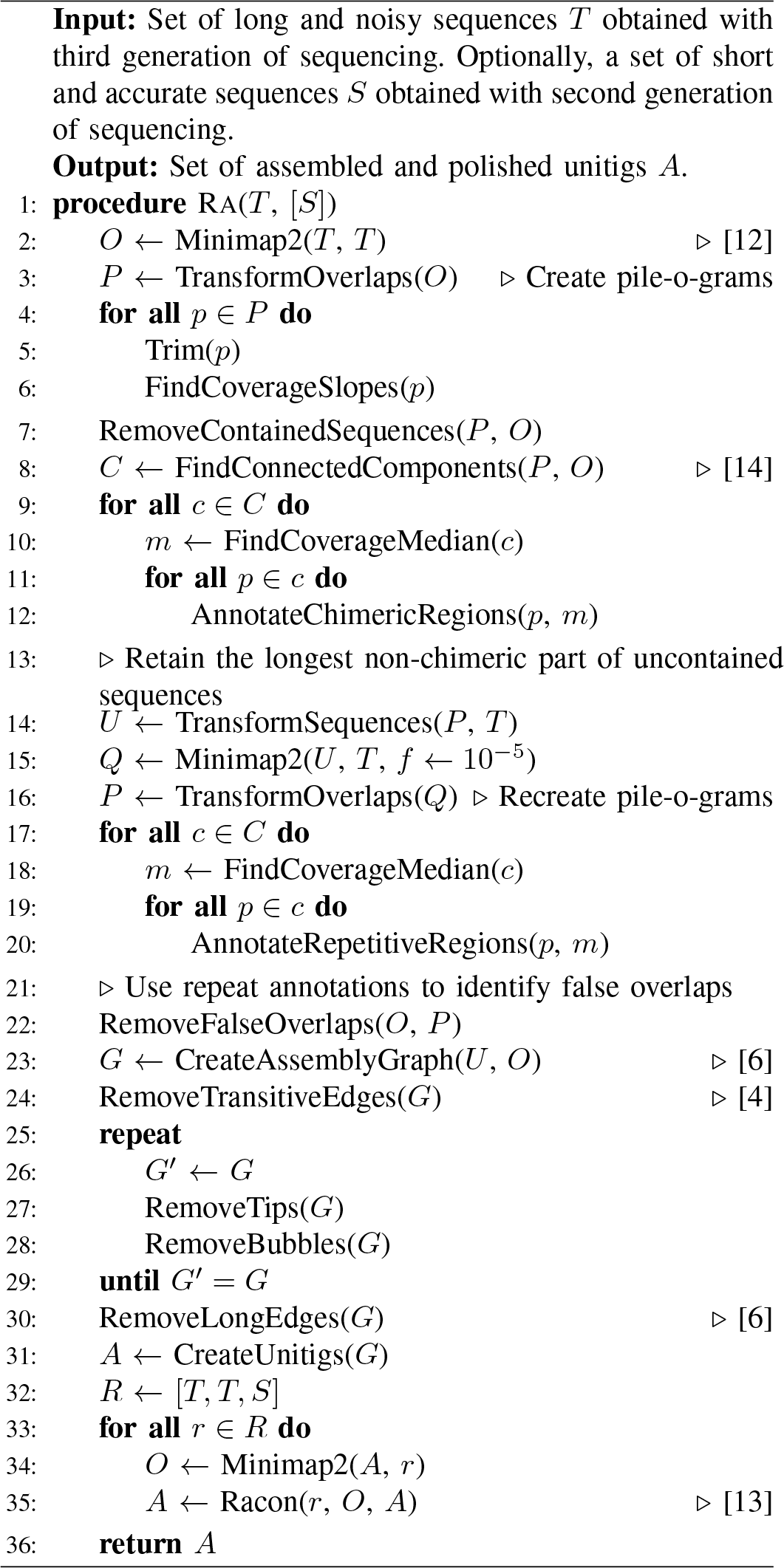
Ra algorithm for de novo genome assembly.

### A. Preprocessing

Pile-o-grams are an impressive tool to dive deeper into the assembly problem, i.e. identification of sequence types. They can be created by stacking all pairwise overlaps of a sequence on top of each other. Summing up the number of overlaps covering each base yields a one-dimensional signal that has a characteristic outline depending on the sequence type. Sequences that uniquely and fully map to the sequenced genome should have almost uniform coverage across their length as shown in Fig. 1. Others have noticeable fluctuations in the signal. Chimeric sequences are sequencing artefacts consisting of multiple parts which are arranged in a way that is absent in the genome. Such signals mostly have a sharp decline in coverage which is followed with a sharp increase, between each of the connected parts (Fig. 2a). There are also cases when those parts overlap a certain amount, causing a spike that reminds of a Dirac delta function (Fig. 2b). Sequences containing a repetitive region at either of the ends can spawn false suffix-prefix overlaps. Those regions can be identified as an increase in coverage in form of a step function (Fig. 2c).

**Fig. 1.**
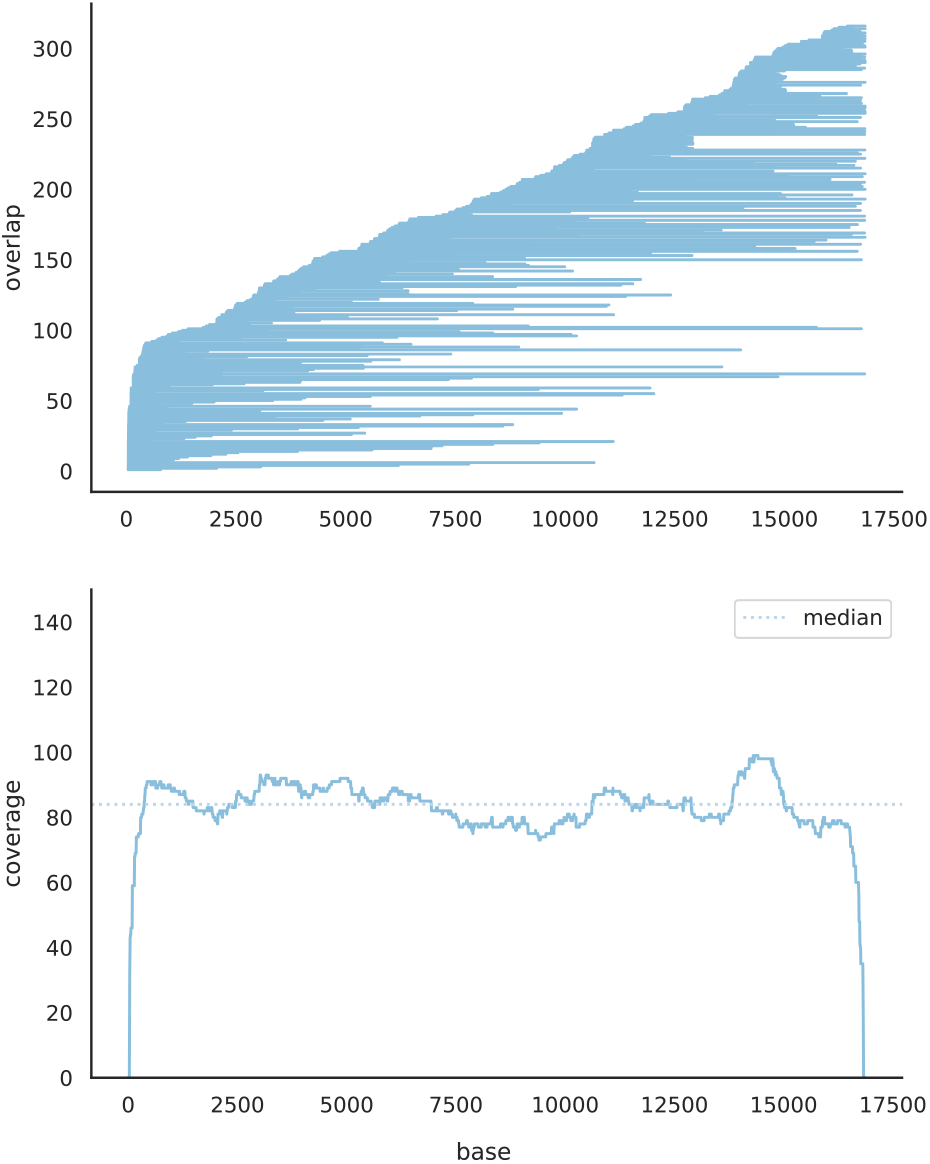
Pile-o-gram that is almost uniform. It was obtained by adding up the number of overlaps covering each base in a sequence which can be uniquely mapped to the reference genome.

**Fig. 2.**
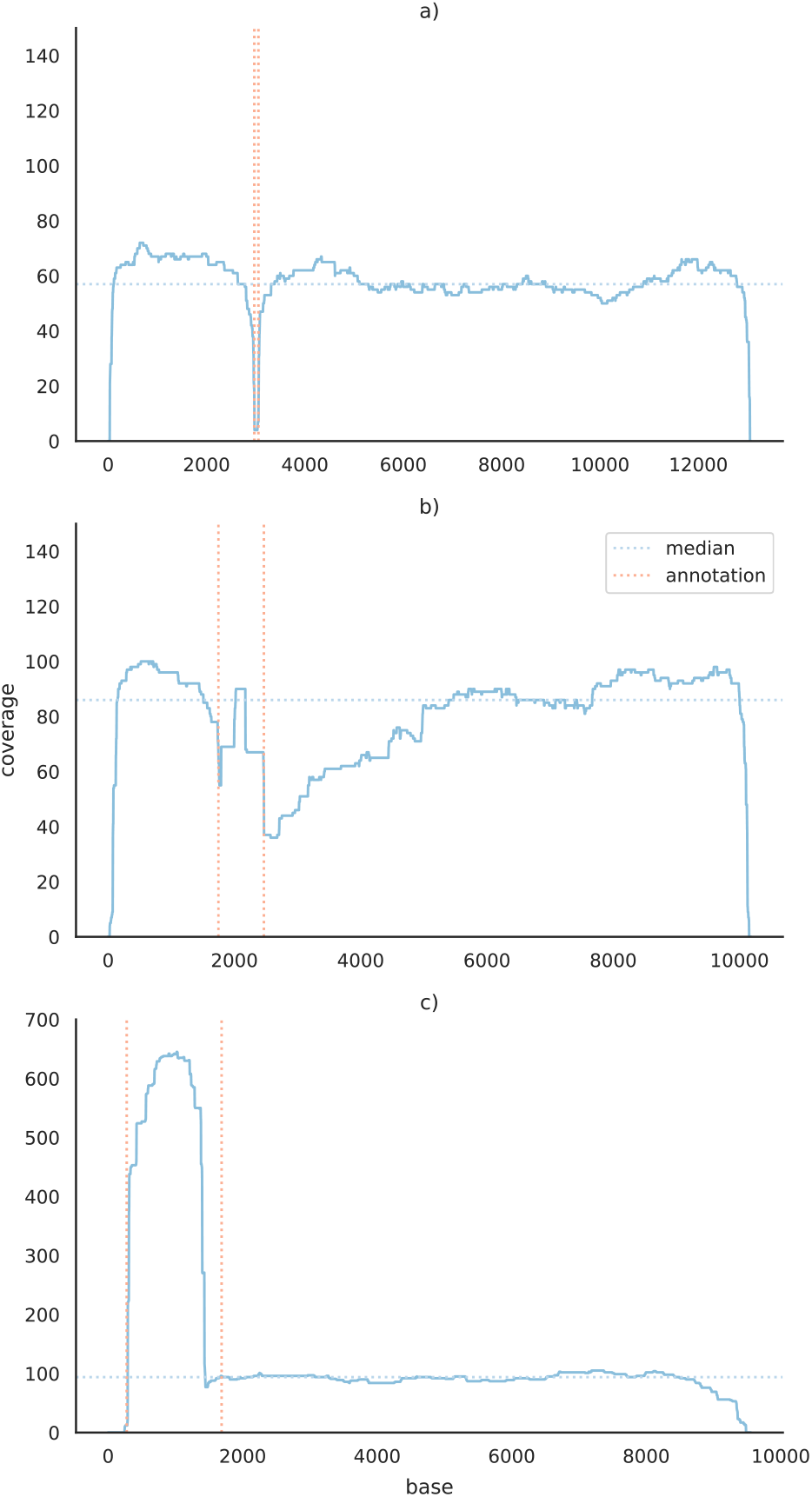
Pile-o-gram representatives for other sequence types with highlighted regions of interest. Chimeric sequences, those that partially map to different regions of the genome, can be seen in subfigures a) and b). The pile-o-gram in a) has a rift denoting that there is almost no overlaps covering this part of the sequence. The one in b) has a narrow ridge that indicates a little overlapping region between parts that constitute this chimeric sequence. On the other hand, repetitive regions in pile-o-grams can be identified by large ridges as seen in pile-o-gram c).

Hinge uses pile-o-grams to detect chimeric sequences and annotate repetitive genomic regions by calculating coverage gradients [10]. Chimeric sequences are truncated to the longest non-chimeric part. On the other hand, repeat annotations are used to find sequences that do not fully bridge certain repeats to allow some of them multiple overlaps in an otherwise best overlap graph, and later use this information for repeat resolution [10].

In the first stage Ra implements a similar approach. All pairwise overlaps are loaded in blocks and immediately transformed into pile-o-grams to decrease the memory footprint.

At this point, each sequence is represented with a vector of unsigned short integers containing base coverages which is approximately equal to storing the complete FASTQ version of the input sequence file. Such pile-o-grams are first used to trim sequences to the largest region covered by at least three sequences, similarly to Miniasm [6]. Afterwards, Ra scans through the coverage vector to identify slopes in the signal by keeping sliding windows left and right of each position in the sequence. The maximal value of each window is compared to the current position and the slope is stored if the coverage ratio is large enough. Collected slopes are grouped into more complex shapes such as rifts and ridges, and each group is further investigated. To decrease the number of false positive annotations, we utilize the information about approximate sequencing depth. The pairwise overlaps are parsed again, contained sequences are dropped, internal overlaps are set aside, and only the remaining prefix-suffix overlaps are kept. They are used to group sequences in connected components with a simple depth-first search [14]. Coverage medians are calculated from pile-o-grams and a global median is obtained for each connected component separately. This way we can treat different sequencing depths correctly. The coverage median of a component is used to determine the relevance of each rift and ridge in all signals of that component. A rift is chimeric if it contains a base with coverage bellow the coverage median divided by 1.84, while ridges represent repetitive regions if majority of their bases have coverage above the coverage median multiplied with 1.42 (both values empirically determined). Narrow ridges located inside the middle of each sequence also undergo a chimeric test. During the second parsing of the overlap file they are declared chimeric if there are not at least three overlaps containing them. Once annotation is finished, Ra breaks sequences over rifts and chimeric ridges, retains the longest non-chimeric region and reconsiders corresponding overlaps that have been classified as internal beforehand. Remaining overlaps that either start or end inside a ridge located at a sequence end are removed, if at least one of overlapped sequences has another overlap that pierces through the ridge in question.

Ra was developed atop minimap2 because it is the fastest pairwise overlapper for long reads. Minimap2 and its predecessor minimap both filter out the most frequent k-mers in order to decrease the number of matching minimizers bound to be stored into memory and thus increases the execution speed [6] [12]. The affected k-mers mostly originate from repetitive regions or from higher copy-number molecules (e.g. plasmids). By employing the layout step hierarchically, first removing contained and chimeric sequences in the preconstruction step, we postpone the repeat annotation to the second step. The whole sequence set is mapped only against the leftover sequences (which constitute a tiny bit of the dataset) and the k-mer filter is decreased by at least one order of magnitude. Pile-o-grams are constructed anew and the increase in base coverage of repetitive regions greatly aids the annotation process.

As mentioned before, pile-o-grams can be treated as one-dimensional signals which makes them suitable for machine learning algorithms. Therefore, we also applied semi-supervised [15] and unsupervised learning algorithms [16] in order to aid sequence classification and annotation. Unfortunately, current results are still outperformed by heuristics.

### B. Graph simplification

After preprocessing, the assembly graph is built and simplified stepwise. Transitive reduction is applied first, as described in [4]. Next, paths consisting of less than seven sequences and without any incoming edges are treated as dead ends in the graph (tips) and are removed in an iterative fashion. Finally, bubble-like structures are found with breadth-first search similar to [17]. Each bubble is inspected for edges whose removal will not cause a break in any other path of the graph (Fig. 3). The path of the bubble with fewer sequences is examined first, and if such edges do not exist, the other path is considered. Bubbles are detected and popped iteratively as well.

**Fig. 3.**
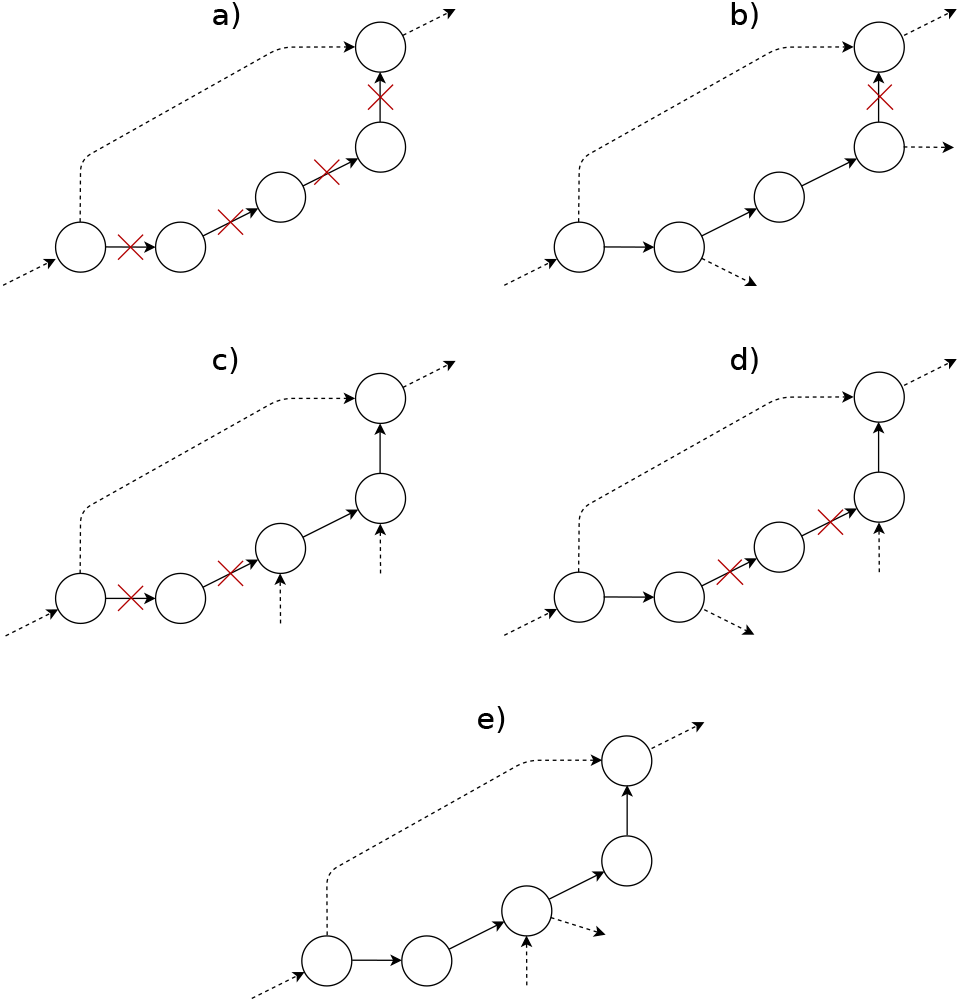
Path types in bubble-like structures of the assembly graph. Paths in consideration consist of solid lines and are inspected for edges which removal will not discontinue any other path. Subfigure a) depicts the simplest scenario in which the whole path can be removed. When they are several nodes with multiple outgoing edges and a single incoming edge, everything after the last node can be deleted as in b). Similar rule applies to the case in c) where they are several nodes with multiple incoming edges and a single outgoing edge. Everything before the first such node can be deleted. When there is a combination of those node types, the edges after the last node with multiple outgoing edges and before the first node with multiple incoming edges can be deleted, as seen in d). This only applies when those two nodes do not have multiple edges of the other type, i.e. if a node has multiple incoming edges it may not have multiple outgoing edges (subfigure e)).

Above described simplification methods coupled with sequence preprocessing are often enough to fully reconstruct the genome. Sequence preprocessing being a heuristic method can sometimes miss a portion of chimeric sequences and false overlaps, leaving us with tangles in the assembly graph. Miniasm solves its leftover tangles by removing short overlaps [6]. Ra follows the same approach but removes only those overlaps that are much shorter than any other overlap in a given tangle (4.2 times by default).

## III. RESULTS

We evaluated our de novo assembler on several publicly available Oxford Nanopore and Pacific Biosciences data sets. Results such as NG50, memory consumption and CPU time can be seen in Table II. We run Ra (commit *07364a1*) with 12 threads on a machine with Intel Xeon CPU (2.40GHz), and used QuastLG [18] (*v5.0.2*) and dnadiff [19] (*v1.3*) for evaluation.

**TABLE II.** ASSEMBLY EVALUATION

| Dataset | Species | Type | Size<br>($\times 10^6$) | Depth | Length | NG50 | Identity | CPU time (min) / Memory (GB) | | |
| --- | --- | --- | --- | --- | --- | --- | --- | --- | --- | --- |
|  |  |  |  |  |  |  |  | Overlap step | Layout step | Consensus step |
| SRR5665597 | *Klebsiella pneumoniae* | ONT | 5.3 | 122 | 5453526 | 5344601 | 0.9841 | 26.39 / 8.32 | 3.71 / 3.19 | 116.45 / 2.22 |
| ERR1140973 | *Klebsiella pneumoniae* | PB | 5.3 | 45 | 5548584 | 5299089 | 0.9915 | 4.20 / 3.45 | 1.10 / 1.02 | 22.04 / 0.79 |
| SCONT* | *Saccharomyces cerevisiae* | ONT | 12.1 | 59 | 12166217 | 288671 | 0.9740 | 36.01 / 8.62 | 4.49 / 3.03 | 73.53 / 2.20 |
| SCPB** | *Saccharomyces cerevisiae* | PB | 12.1 | 127 | 12267357 | 711780 | 0.9974 | 47.03 / 16.87 | 8.20 / 4.54 | 122.89 / 4.702 |
| SRR6702603 | *Drosophila melanogaster* | ONT | 143.7 | 32 | 128449879 | 1854698 | 0.9884 | 652.06 / 22.94 | 53.35 / 10.82 | 570.28 / 13.75 |
| SRX499318 | *Drosophila melanogaster* | PB | 143.7 | 109 | 135936009 | 2128249 | 0.9976 | 2302.29 / 34.15 | 676.85 / 51.92 | 1669.46 / 47.45 |

\*S288C R9 dataset from http://www.genoscope.cns.fr/externe/Download/Projets/yeast/datasets/raw_data/
\*\*W303 P4C2 dataset from https://github.com/PacificBiosciences/DevNet/wiki/Saccharomyces-cerevisiae-W303-Assembly-Contigs

We can observe how NG50 can still be improved and we believe that this is achievable with more relaxed constraints in sequence annotation and better heuristics for solving leftover tangles in the assembly graph. The memory consumption is bound by the sequence file plus some overhead for the overlap files which should be small enough for most use cases. The overlap and consensus steps dominate the execution time, but we believe it is possible to speed up the overlap step, which is the future path of optimizations we will consider.

Ra is a simple and lightweight assembler which happens to perform well on genomes sizes ranging from bacteria to plants shown by independent evaluations. For example, in a comparison of long read assembler applied to various bacterial datasets [20]), Ra was declared the most reliable assembler. In addition, it was used by [21] and yielded the most contiguous plant assemblies in their study.

